# Engineered post-transcriptional regulation of *Scn10a* by Pumilio ameliorates neuropathic pain in mice

**DOI:** 10.64898/2026.11.13.765935

**Authors:** Guadalupe García, Jacob B. Shapiro, Alexander R. Mikesell, James P. Falese, Kai A. Bartol, Katherine M. McKenney, Shea Ruhly, Aaron C. Goldstrohm, Robert A. Pearce, Traci M. Tanaka Hall, Zachary T. Campbell

## Abstract

Post-transcriptional control of mRNA permeates biology, yet its contribution to sensory neuron physiology and pain remains unclear. The mammalian Pumilio proteins PUM1 and PUM2 (collectively referred to as PUM) are RNA-binding proteins that bind Pumilio response elements (PREs), typically situated in 3 ′ untranslated regions. PUM recruits the CCR4-NOT complex to promote mRNA deadenylation and decay. Several mRNAs that encode voltage-gated sodium channels that regulate action potential generation in pain sensing neurons harbor PREs. Here, we explored whether PUM plays a role in pain signaling. PUM1 and PUM2 were enriched in TRPV1-positive dorsal root ganglion neurons, where PUM2 physically interacted with *Scn9a*, *Scn10a*, and *Scn11a* transcripts, which encode Na_v_1.7, Na_v_1.8, and Na_v_1.9, respectively. Depletion of PUM in sensory neurons produced mechanical hypersensitivity and enhanced inflammatory pain. These effects were reversed by blocking Na_v_1.8 but not Na_v_1.7, raising the possibility that repression of *Scn10a* by PUM can be harnessed using synthetic biology to diminish neuropathic pain. To this end, we generated a chimera comprising the RNA-binding domain of PUM2 and a repressive domain, which we dubbed ‘super-PUM’. Expression of AAV-encoded super-PUM in sensory neurons reduced Na_v_1.8 abundance, diminished tetrodotoxin-resistant and suzetrigine-sensitive sodium currents, and prevented mechanical and thermal hypersensitivity following spared nerve injury. These results demonstrate that optimization of an endogenous post-transcriptional regulon provides a means to ameliorate neuropathic pain.

## INTRODUCTION

Post-transcriptional control of mRNA is pervasive (*1–4*). mRNAs harbor two untranslated regions (UTRs) situated at the 5′ and 3′ ends of the transcript. Information present within the 3′ UTR governs every aspect of mRNA function including export, subcellular localization, translation, and stability. Regulatory elements in UTRs are encoded by cis-acting elements and decoded by trans-acting elements such as RNA-binding proteins (RBPs) and non-coding RNAs. While the biological importance of regulation encoded within the 3′ UTR is incontrovertible, little is known regarding its roles in nociceptive sensory neurons that detect and amplify painful cues.

PUM1 and PUM2 are archetypal post-transcriptional regulatory factors. Mammals express these two paralogous PUM genes, which encode RNA-binding domains with 91% amino acid identity (*5–7*). PUM binds to a highly conserved 8-nucleotide consensus element (UGUANAUA), known as the Pumilio response element (PRE) (*8–11*). Once bound to a target mRNA, PUM recruits the CCR4-NOT deadenylase complex through an intrinsically disordered N-terminal domain (*12–14*). Assembly of the repressive complex promotes deadenylation, translational repression, and ultimately mRNA decay. In the mammalian CNS, PUM regulates synaptic plasticity, neurodevelopment, and dendrite formation (*15–17*). However, it is unclear what, if any, role PUM play in the PNS.

Voltage-gated sodium channels (VGSCs) play pivotal roles in pain (*18*). Na_v_1.7, Na_v_1.8, and Na_v_1.9, (encoded by *Scn9a*, *Scn10a*, and *Scn11a*, respectively) are integral to action potential generation and propagation (*18, 19*). Gain-of-function mutations in each factor are associated with hypersensitivity to sensory inputs in humans, whereas loss-of-function mutations are linked to pain insensitivity (*18, 20*). In *Drosophila*, Pumilio directly regulates the VGSC Para, impacting motor neuron excitability (*21*). More recently, Pumilio was also shown to regulate nociceptive sensitivity in larvae (*22*). However, whether PUM proteins regulate either the stability or translation of mRNAs that encode VGSC transcripts in mammalian nociceptors remains unclear.

Here, we show that PUM1 and PUM2 are preferentially expressed in TRPV1-positive, peptidergic sensory neurons in DRG tissue. Immunoprecipitation of PUM2 revealed physical association with *Scn9a*, *Scn10a*, and *Scn11a* transcripts. *In vivo,* CRISPR-Cas9-mediated depletion of PUM in sensory neurons resulted in an increase in Na_v_1.7, Na_v_1.8, and Na_v_1.9. PUM depleted mice displayed mechanical allodynia which was reversed by pharmacologic blockade of Na_v_1.8. To examine the effects of PUM supplementation, we engineered a hyper-repressive chimera, termed super-PUM, consisting of the PUM2 RNA-binding domain fused to the CCR4-NOT deadenylase subunit CNOT7, which represses translation when tethered directly (*23*). AAV-mediated delivery of this construct to sensory neurons suppressed mechanical and thermal hypersensitivity caused by endogenous PUM loss. Introduction of super-PUM resulted in significant reduction of Na_v_1.8 levels in DRG tissue and reduction of TTX-resistant and suzetrigine-sensitive sodium currents. Finally, super-PUM prevented the development of mechanical and thermal hypersensitivity in the spared nerve injury (SNI) model of neuropathic pain. Together, our findings reveal coordinate regulation of VGSCs by PUM and demonstrate that ectopic delivery of PUM is a potential strategy for alleviation of neuropathic pain.

## RESULTS

### PUM2 associates with mRNAs that encode VGSCs

To explore if PUM is expressed in sensory neurons, we examined murine DRG tissues using immunohistochemistry (**Fig. 1A-B**). PUM levels were quantified in peptidergic (TRPV1+) and non-peptidergic (IB4+) nociceptors. In these cells, PUM1 was predominantly expressed in the TRPV1^+^ population (55%). Approximately 10% of IB4^+^ neurons expressed PUM1 (**Fig.1A**). Similarly, PUM2 was more abundant in TRPV1^+^ neurons, where it was expressed in nearly 50% of cells, while it was present in less than 25% IB4^+^ neurons (**Fig.1B**). Next, we tested if a subset of mRNAs were physically associated with PUM2 using RNA immunoprecipitation (RIP) from lumbar DRGs, followed by RT-qPCR (**Fig. 1C**). PUM2 capture was effective based on immunoblot analysis of immunoprecipitant (**Suppl. Fig. 1**). Two previously identified PUM2 targets, *Megf9* and *Gsk3b*, were highly enriched following PUM2 immunoprecipitation (*24*) (**Fig. 1 D-E**). Importantly, the negative control *Elavl1*, which lacks PREs, was not enriched (**Fig. 1F**). PUM2 association was quantified for all VGSCs. PUM has been linked to Na_v_1.6 in rat cortical neurons (*25*). We did not detect a direct association potentially indicative of tissue-specific patterns on sodium channel mRNA expression. However, transcripts encoding *Scn9a* (Na_v_1.7), *Scn10a* (Na_v_1.8), and *Scn11a* (Na_v_1.9), were significantly enriched (**Fig. 1G-P**). Importantly, all three gene products have been linked to pain in humans and in rodents (*18*). This observation prompted us to investigate if PUM plays a role in pain signaling.

**Figure 1.**
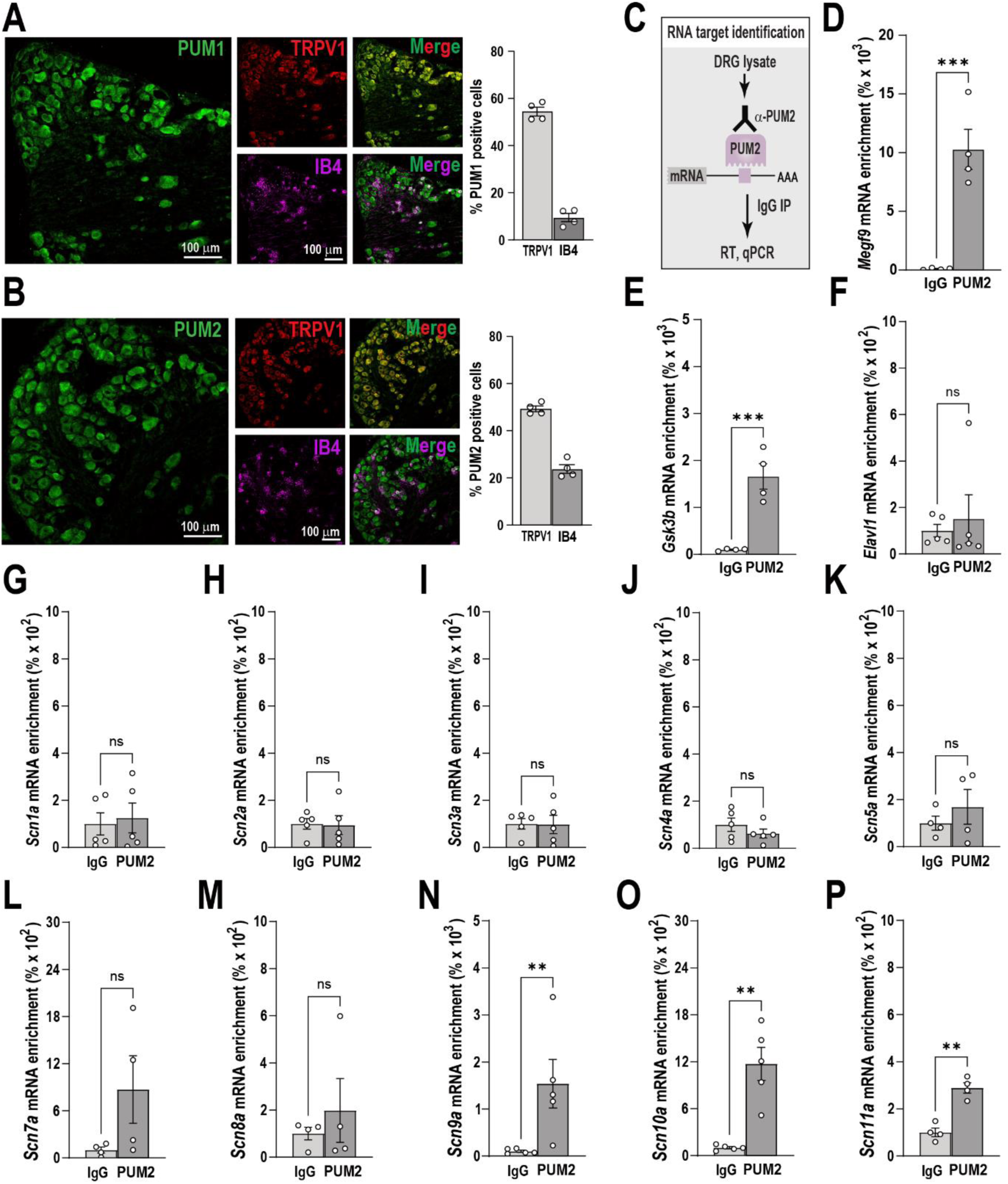
PUM expression and interaction with mRNAs that encode voltage gated sodium channels in DRG tissues. (**A**) PUM1 and (**B**) PUM2 expression in lumbar dorsal root ganglia (DRG) sections from wild-type mice. PUM1 was predominantly expressed in TRPV1^+^ neurons (colocalization rate of 54.71 ± 3.39%), whereas its expression in IB4^+^ neurons was considerably lower (9.48 ± 3.00%). Likewise, PUM2 was more frequently detected in peptidergic (TRPV1+) neurons (49.33 ± 2.08%) than in non-peptidergic (IB4+) neurons (23.80 ± 3.23%). Bars represent the mean ± SEM of relative intensity values from *n* = 4 mice per group. Scale bar: 100 µm. (**C**) Scheme of RNA immunoaffinity purification (RIP) of PUM2 from DRG lysates obtained from wild-type mice. (**D-P**) RIP was performed using antibodies against PUM2 or IgG, followed by quantification of target transcripts by qPCR. Luciferase mRNA was used to normalize mRNA levels of the positive controls *Megf9* and *Gsk3b*, the negative control *Elav1*, and all voltage-gated sodium channels family (*Scn1a*-*Scn11a*) transcripts. Bars represent the mean ± SEM of 4-5 mice per group, showing the percent of mRNA enrichment between samples treated with PUM2 or IgG antibodies, calculated using the 2^-(ΔΔCт) method. **P < 0.01, ***P <0.001, ns (not significant), one-way ANOVA followed by Bonferroni’s multiple comparisons test.

### Genetic depletion of PUM results in mechanical hypersensitivity

To test if PUM is involved in somatosensation, we depleted PUM1 and PUM2 specifically in sensory neurons. We previously reported an efficient *in vivo* genetic perturbation strategy wherein transgenic SpCas9 expression is restricted to sensory neurons and guide RNAs are injected into the intrathecal space (**Fig. 2A**) (*26*). This approach enables dispersion of guide RNAs bilaterally to lumbar DRGs and achieves up to 65% target reduction *in vivo*. We simultaneously depleted PUM1 and PUM2 by delivering pooled guide RNAs targeting both genes. To quantify PUM depletion, protein levels were assessed using immunohistochemistry on DRG tissue slices (**Fig. 2B-C**). For both PUM1 and PUM2, greater than 50% depletion was observed. To determine if PUM regulates the abundance of polypeptides encoded by *Scn9a, Scn10a*, or *Scn11a*, immunoblots were conducted on DRG tissues (**Fig. 2D**). Following PUM depletion, all three proteins displayed an increase in abundance of 58%, 64%, and 174% for Na_v_1.7, Na_v_1.8, and Na_v_1.9, respectively. Next, we examined evoked sensitivity to mechanical and thermal cues in PUM-depleted mice. We observed increased baseline mechanical sensitivity in PUM1 and PUM2-depleted male and female mice (**Fig. 2E**). Individual depletion of either PUM1 or PUM2 had no effect on baseline mechanical sensitivity (**Suppl. Fig. 2**), suggesting functional redundancy of these proteins. Injection of a mild inflammatory stimulus, prostaglandin E_2_ (PGE2), in the hindpaw evoked mechanical hypersensitivity in both male and female control and PUM-depleted mice. In contrast, combined PUM depletion did not alter baseline sensitivity to noxious heat but significantly enhanced PGE2-induced heat hyperalgesia compared to controls (**Fig. 2F**). These observations indicate that PUM regulates the abundance of key VGSCs involved in baseline mechanical sensation and inflammatory heat hyperalgesia.

**Figure 2.**
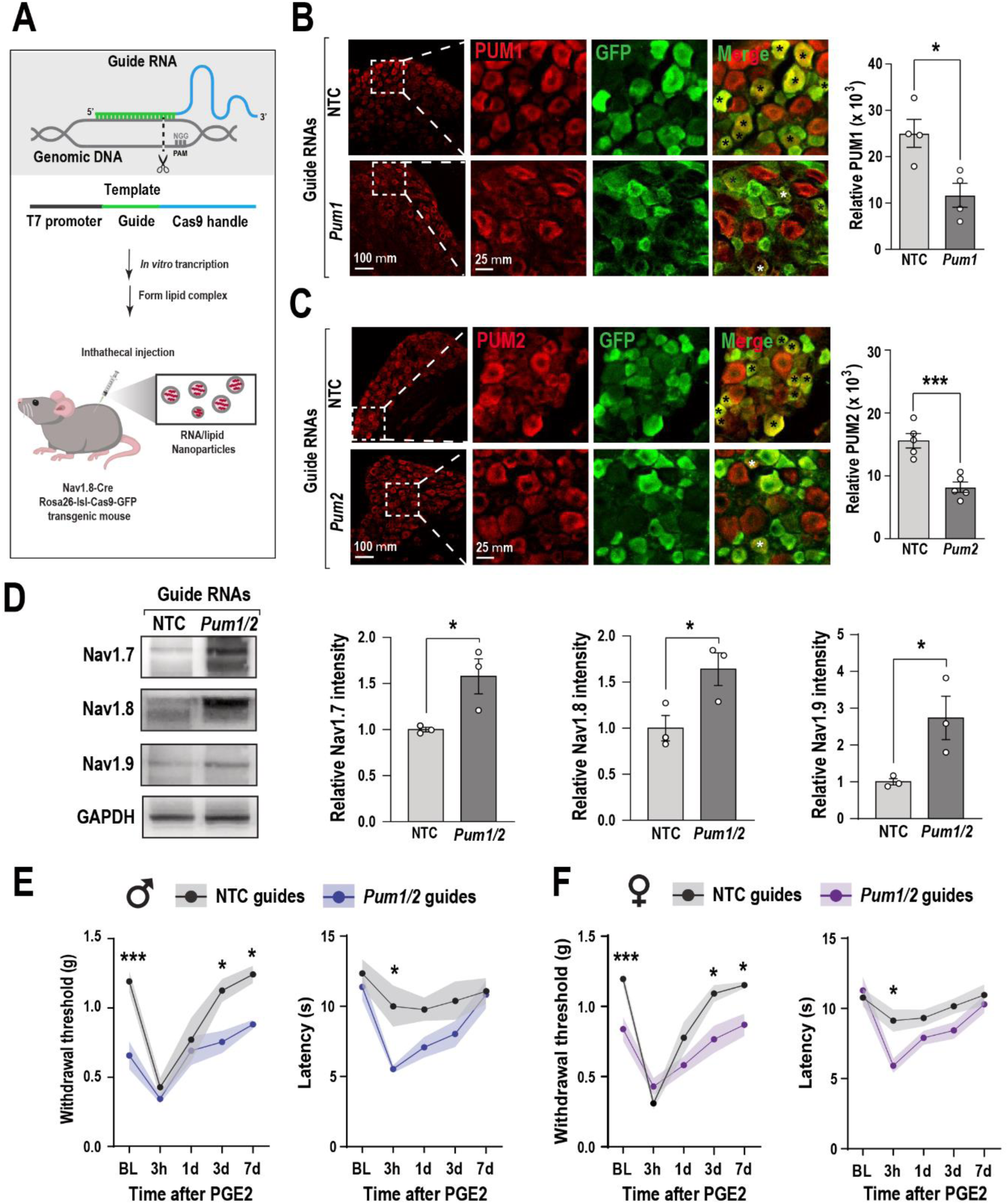
Depletion of PUM1 and PUM2 in mice influences sodium channel abundance and baseline responses to mechanical stimulation. (**A**) Scheme of RNA guide synthesis and *in vivo* delivery. The DNA template for RNA guide synthesis includes a T7 promoter, the gene-specific sgRNA sequence, and elements essential for Cas9 enzyme function. RNA guides were produced by *in vitro* transcription and RNA-lipid complexes were prepared for intrathecal delivery into Cas9 knock-in mice. In these animals, SpCas9 and GFP are expressed from the same transgene (*53*). (**B**) Immunohistochemical staining of PUM1 or (**C**) PUM2 expression in DRG sections after two intrathecal injections of 3 µg *Pum1/Pum2* guide RNA pool or NTC (non-targeting control). The relative intensity of PUM1 or PUM2 expression was quantified only in Cas9^+^ (GFP^+^) cells. PUM1 and PUM2 expression were reduced in *Pum*-guide injected mice compared to NTC. Bars represent the mean ± SEM of relative intensity values from n = 4-5 mice per group. *P < 0.05 or ***P < 0.001 was determined by an unpaired t-test. Scale bar: 100 µm for whole DRG images and 25 µm for magnified views. (**D**) Representative blots showing the relative abundance of Na_v_1.7, Na_v_1.8 and Na_v_1.9 protein expression obtained from DRGs of Cas9 knock-in mice treated with *Pum* or NTC guides. Bars represent the mean ± SEM of relative protein abundance normalized to GAPDH from 3 mice per group. *P < 0.05 was determined by an unpaired t-test. Male (**E**) or female (**F**) mice were assessed following intraplantar delivery of PGE_2_ (100 ng) for mechanical and thermal hypersensitivity after injection of *Pum* or NTC guides. Lines represent the mean ± SEM (shaded area) of withdrawal threshold (g) or latency (s) from 6-8 mice per group. Measurements were performed before PGE_2_ injection (baseline, BL) and at 3 hours (3h), 1 day (1d), 3 days (3d) and 7 days (7d) post-injection. *P < 0.05, ***P < 0.001 were determined by two-way ANOVA followed by Bonferroni’s post hoc test.

### PUM PREs in *Scn9a* confer regulation by PUM

The PUM consensus binding motif is UGUANAUA (*8, 11, 27*). We identified five putative PREs in the *Scn9a* 3′UTR and confirmed their association with recombinant PUM2 using electrophoretic mobility shift assays (EMSAs) (**Suppl. Fig. 3**). All the equilibrium dissociation values (*K_d_)* ranged from 36.0 – 90.5 nM. This range is comparable to known PUM targets (*5*). Mutation of the 5′ UGU element to ACA abrogates binding and regulation by PUM and its homologs (*9, 28*). To determine if regulation of *Scn9a* by PUM is functionally important *in vivo*, we generated a strain in which all five of the putative regulatory elements were mutated from UGU to ACA (designated *Scn9a^PRE-Mut^*) (**Fig. 3A**). Loss of the five PREs resulted in an increase in Na_v_1.7 specifically in TRPV1 expressing sensory neurons (**Fig. 3B-C**). To confirm that PUM2 association with other mRNA targets was preserved, we performed RNA immunoprecipitation on DRGs isolated from *Scn9a^PRE-Mut^* animals. The PUM2 target *Megf9* was enriched, while the negative control *Elavl1* was not (**Fig. 3D-E**). PUM2 bound to *Scn10a* and *Scn11a* in mutant samples while association with *Scn9a* was lost (**Fig. 3F-H Fig. 1N**). Together, these findings support direct recognition and regulation of *Scn9a* by PUM.

**Figure 3.**
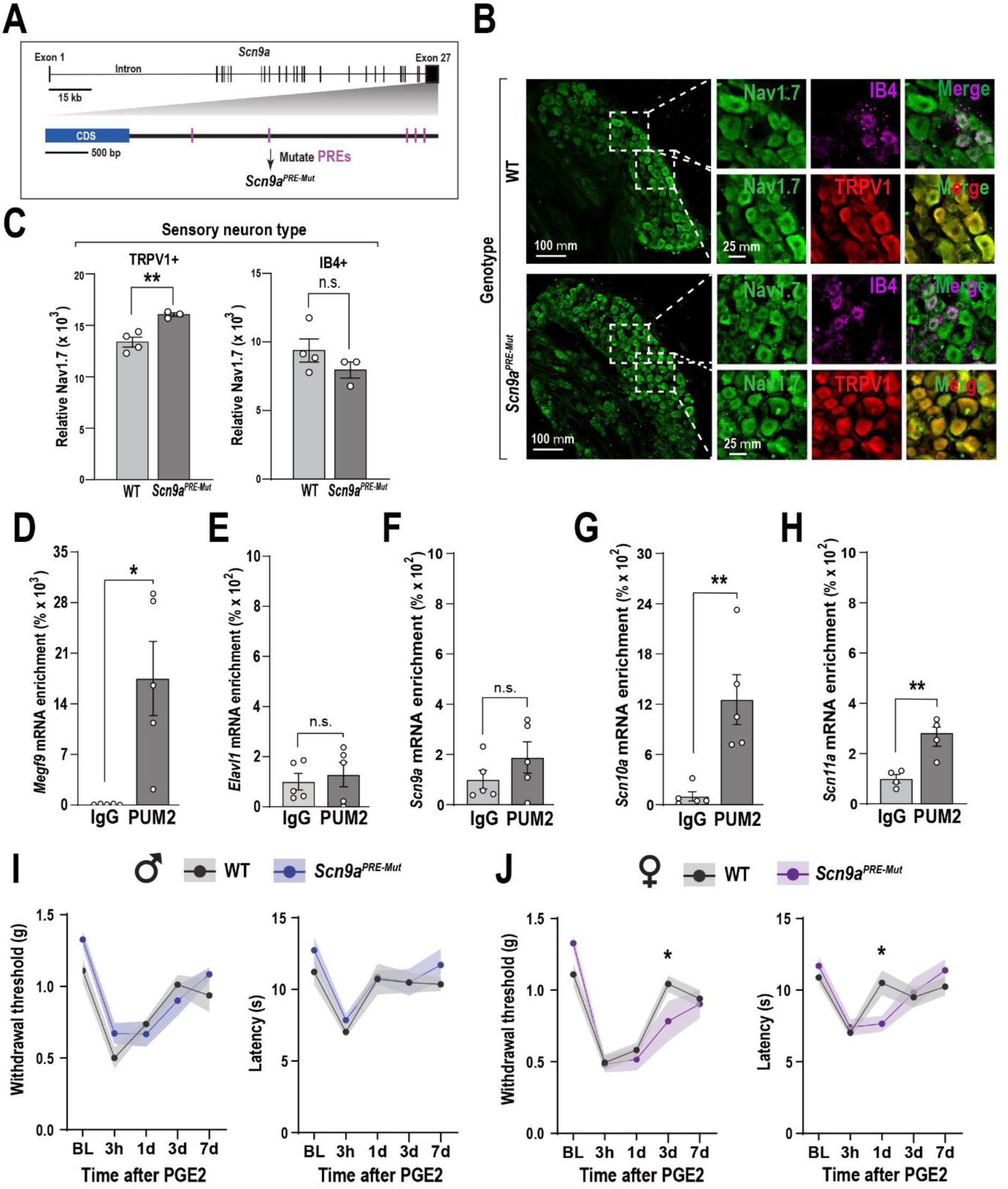
Loss of PREs in *Scn9a* confers minimal changes in mechanical and thermal responses to PGE2. (**A**) A schematic of the *Scn9a*gene showing the location of predicted Pumilio response element (PRE) sites. Five identified PREs were mutated to generate *Scn9a^PRE-Mut^* mice. (**B**) Immunohistochemical staining of Na_v_1.7 expression in DRG sections from WT and *Scn9a^PRE-Mut^* mice. Scale bar: 100 µm for whole DRG images and 25 µm for magnified views. (**C**) Relative Na_v_1.7 intensity was quantified in TRPV1+ and IB4+ neurons. A significant increase in Na_v_1.7 expression was observed in TRPV1+ neurons from *Scn9a^PRE-Mut^* mice, but not in IB4+ cells. Bars represent the mean ± SEM of relative intensity values from n = 3-4 mice per group. **P < 0.01 was determined by an unpaired t-test. (**D-H**) RT-qPCR analysis of DRG lysates obtained from WT and *Scn9a^PRE-Mut^* mice following RIP with anti-Pum2 or IgG antibodies. Luciferase mRNA was used for normalization. Enrichment of *Megf9* (positive control), *Elavl1* (negative control), and the *Scn9a*, *Scn10a* and *Scn11a* transcripts are shown. Bars represent the mean ± SEM of 4-5 mice per group, showing the percent of mRNA enrichment between samples treated with PUM2 or IgG antibodies, calculated using the 2^-(ΔΔCт) method. *P < 0.05, **P <0.01, n.s. (not significant), after one-way ANOVA followed by Bonferroni’s multiple comparisons test. (**I-J**) Mechanical and thermal hypersensitivity were assessed following intraplantar delivery of PGE_2_ (100 ng) in male (**I**) and female (**J**) *Scn9a^PRE-Mut^* mice. Lines represent the mean ± SEM (shaded area) of withdrawal threshold (g) or latency (s) from 6-8 mice per group. Measurements were taken at baseline (BL) and at 3 hours (3h), 1 day (1d), 3 days (3d) and 7 days (7d) post-injection. *P < 0.05, was determined by two-way ANOVA followed by Bonferroni’s post hoc test.

Next, behavioral responses to PGE_2_ were quantified. Baseline sensitivities in the *Scn9a^PRE-Mut^* strain were not different than wild-type mice (**Fig. 3I-J**). A small increase in mechanical and thermal responsiveness at the one-day timepoint was observed specifically in females. However, this change was modest and transient. We conclude that PREs situated in the 3′ UTR of *Scn9a* confer regulation in sensory neurons. However, the lack of substantial changes in pain-associated behaviors in *Scn9a^PRE-Mut^* mice indicates that phenotypes observed in PUM depleted animals are unlikely to be driven by Na_v_1.7.

### Inhibition of Na_v_1.8 reverses mechanical allodynia in PUM depleted mice

Given the subtle phenotype observed in the *Scn9a^PRE-Mut^* strain, we asked whether pharmacologic blockade of either Na_v_1.8 or Na_v_1.7 reversed mechanical hypersensitivity in PUM depleted animals. We focused on mechanical allodynia in these mice given that this phenotype is robust and not sexually dimorphic. Suzetrigine (Journavx) is a selective inhibitor of Na_v_1.8 that is approved for the treatment of acute pain in humans (*29*). Injection of suzetrigine in the hindpaw had no effect on animals that received non-targeting guide RNAs but resulted in a transient but substantial reversal of mechanical hypersensitivity in the PUM-depleted mice (**Fig. 4A**). In contrast, blockade of Na_v_1.7 with the selective inhibitor PF-05198007 had no effect in either group (**Fig. 4B**), consistent with the phenotype of the *Scn9a^PRE-Mut^* strain (*30*). The data indicate that Na_v_1.8 plays a substantial role in mechanical allodynia following PUM depletion. To determine if PUM recognizes PREs situated in *Scn10a*, we quantified binding of recombinant PUM2 to putative binding elements using EMSAs for four putative PREs (**Suppl. Fig. 4**). A single PRE bound with high affinity (*K*_d_ = 82 nM). Two displayed modest to weak binding (*K*_d_ = 167 and 500 nM) and one element did not bind (*K*_d_ > 2,000 nM). We conclude that mechanical allodynia triggered by loss of PUM is reversed upon inhibition of Na_v_1.8 and that PREs situated in the *Scn10a* 3′ UTR associate with PUM2 *in vitro*.

**Figure 4.**
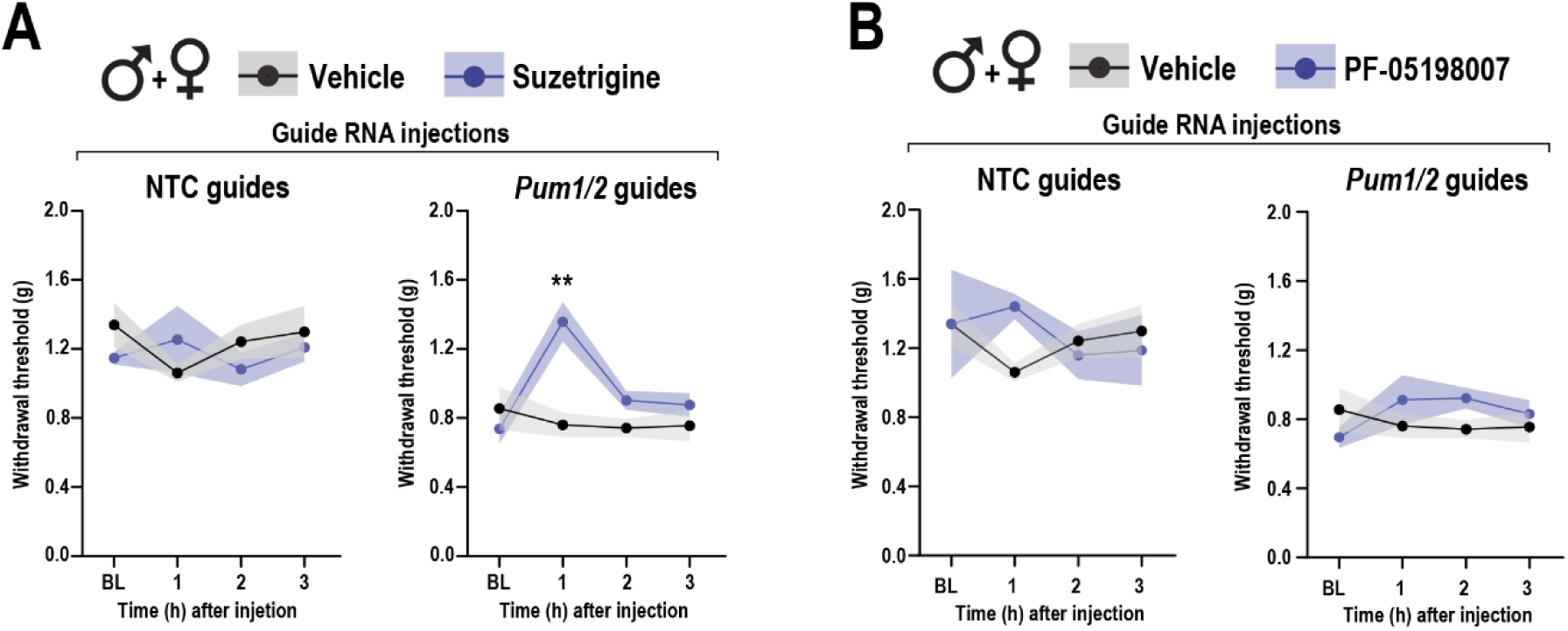
Differential effects of sodium channel inhibitors on mechanical thresholds in PUM-depleted mice. Mechanical hypersensitivity was assessed following intraplantar delivery of suzetrigine (1 µg), PF-05198007 (1 µg), or vehicle in Cas9 mice treated with *Pum1/2* or NTC guides. Lines represent the mean ± SEM (shaded area) of withdrawal threshold (g) from 6-7 mice per group for *Pum1/2* guide injected mice or 3-4 mice per group for NTC controls. Measurements were taken at baseline (BL) and at 1-3 hours after injection. **P < 0.01 was determined by two-way repeated measures ANOVA followed by Bonferroni’s post hoc test.

### Enhancing PUM-mediated repression reduces Na_v_1.8 abundance and function

Identification of Na_v_1.8 as a PUM target prompted us to ask if enhancing PUM mediated repression could further reduce Na_v_1.8 expression and function. PUM2 represses translation through multivalent low-affinity interactions with the CCR4-NOT deadenylase complex (*12*). These interactions are mediated by an extended, N-terminal IDR on PUM2 that encodes several repression domains (*13, 14*). A recent global screen of translational effectors making use of the tethered function assay revealed CNOT7, the deadenylase component of the CCR4-NOT complex, as one of the most repressive polypeptides(*23*). We reasoned that fusion of the RNA-binding domain of PUM2 to CNOT7 would generate a highly repressive version of PUM sufficiently compact to enable ectopic viral expression *in vivo*. In a heterologous reporter assay conducted in HEK293T cell lines, we found that a luciferase reporter containing consensus PUM elements was repressed following overexpression of full length PUM2 (**Fig. 5A**). The PUM2-CNOT7 chimera (henceforth super-PUM) was more repressive than native PUM2, whereas a mutant lacking RNA-binding capability (designated super-PUM mutant: repeat 6 (N923A, Y924A, Q927A) and repeat 7 (S959A, N960A, E963A) showed no repression (*31*).

**Figure 5.**
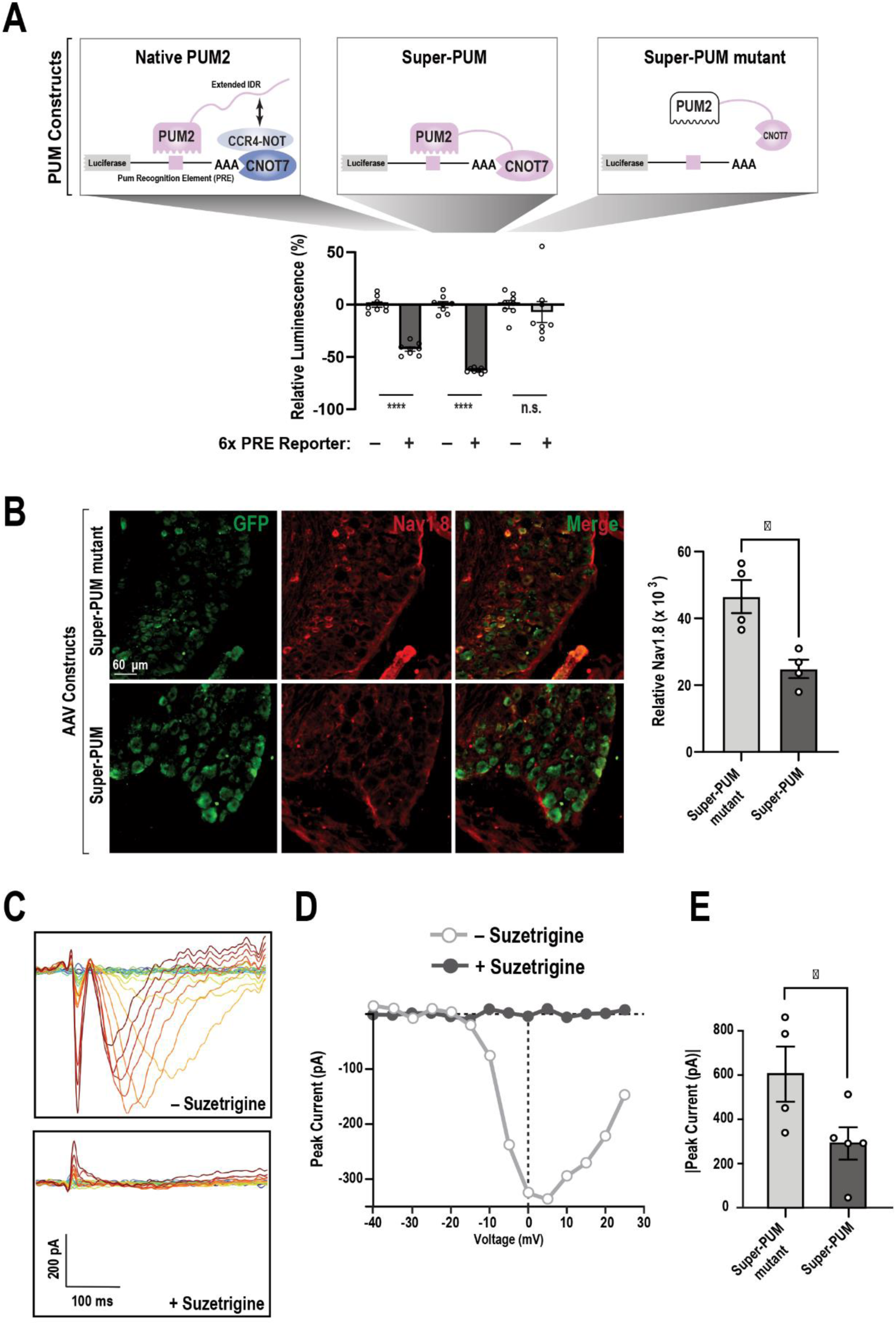
Super-PUM reduces translation of a PRE-containing reporter and diminishes Na_v_1.8 abundance and Suzetrigne-sensitive, TTX-insensitive sodium currents in sensory neurons. (**A**) A schematic representation. The RNA-binding domain of PUM2 was fused to CNOT7, a catalytic component of the CCR4-NOT deadenylase complex, to enhance translational repression of target transcripts. Constructs encoding native PUM2, the engineered PUM2-CNOT7 fusion protein (super-PUM), and an RNA-binding-deficient PUM2 domain (super-PUM mutant) were transfected in HEK293Ts. Repression activity was assessed using a luciferase reporter containing six PREs. Super-PUM exhibited greater repression than native PUM2, whereas the super-PUM mutant failed to repress reporter expression. (**B**) Relative Na_v_1.8 intensity was quantified in GFP^+^ neurons in DRGs collected from mice injected with AAV encoding super-PUM or super-PUM mutant control and a GFP reporter. A significant decrease in Na_v_1.8 expression was observed in super-PUM mice. Bars represent the mean ± SEM of relative intensity values from n = 4 mice per group. *P < 0.05 was determined by an unpaired t-test. **(C)** Whole-cell voltage-clamp traces of TTX-resistant sodium current recorded from a DRG neuron obtained from a mouse injected with super-PUM AAV in the absence (top) and presence (bottom) of suzetrigine. Currents were elicited by a series of voltage steps ranging from -70 to +25 mV in 5 mV increments, 300 ms duration. (**D)** I-V curves of peak TTX-resistant sodium current of the traces from panel C. (**E)** Peak inward TTX-resistant sodium currents from super-PUM mutant and super-PUM groups. Data shown as mean ± SEM (super-PUM mutant; mean: -605, SEM: 125 pA, super-PUM; mean: -291, SEM: 73 pA, *P < 0.05, determined by unpaired t-test). We found no differences in the command voltages that produce peak inward current (super-PUM mutant; median: 7.5 mV, IQR: 13.75, super-PUM; median: 5.0, IQR: 13.75, p > 0.5, Mann-Whitney U).

We next sought to determine whether super-PUM could reduce endogenous Na_v_1.8 expression and function in sensory neurons. We prioritized Na_v_1.8 because its selective inhibition reversed mechanical hypersensitivity following PUM depletion and selective pharmacological tools targeting Na_v_1.9 remain limited (*32*). To enable sensory neuron specific application of our AAV payloads, we used AAV capsid with trophism for DRG sensory neurons (MaCPNS1) (*33*) and a double-floxed and inverted transgene to ensure Cre-dependent expression. To mark AAV-infected neurons, PUM inserts were fused to eGFP with a C-terminal P2A peptide to covalently separate polypeptides. We delivered super-PUM or super-PUM mutant AAVs intrathecally to *Scn10a^Cre/+^* animals which resulted in robust GFP expression in lumbar DRG tissues (**Suppl. Fig. 5**). Immunostaining of lumbar DRGs (L3–L5) near the intrathecal injection site revealed a 46% reduction in Na_v_1.8 intensity in AAV-infected (GFP^+^) neurons from mice injected with AAV encoding super-PUM compared with those receiving AAV encoding the super-PUM mutant control (**Fig. 5B**). To examine if reduced Na_v_1.8 expression translated to altered channel function, we performed whole-cell voltage-clamp recordings from dissociated DRG neurons isolated from mice expressing either super-PUM or super-PUM mutant AAVs. Recordings were restricted to GFP^+^ neurons to ensure analysis of cells expressing the AAV-delivered constructs. Tetrodotoxin (TTX) (500 nM) was applied to block TTX-sensitive VGSCs, allowing isolation of TTX-resistant sodium currents. Depolarizing voltage steps elicited inward currents that were blocked by the selective Na_v_1.8 inhibitor suzetrigine (1 µM) (*34*), indicating the predominant contribution of Na_v_1.8 to the recorded currents (**Fig. 5C-D**). Compared with PUM mutant controls, super-PUM reduced peak current amplitude by 48% (**Fig. 5E**). The command voltages producing the peak inward current did not differ, consistent with a reduction in Na_v_1.8 channel numbers in the super-PUM group, rather than an alteration of channel properties. These observations indicate that synthetic post-transcriptional regulation conferred by viral introduction of super-PUM specifically in DRG sensory neurons disrupts Na_v_1.8 expression and function.

### Ectopic expression of super-PUM diminishes neuropathic pain

Next, we sought to determine whether super-PUM rescues sensory hypersensitivity caused by genetic depletion of PUM. Injection of super-PUM into double PUM knockdown mice increased mechanical withdrawal thresholds compared to the super-PUM mutant control, with a significant effect observed 3 weeks after AAV delivery (**Fig. 6A**). Super-PUM also increased withdrawal latency in the Hargreaves assay, with a significant increase detected 1 week after AAV delivery. These findings indicate that super-PUM confers repression on a PRE-containing transcript and functionally compensates for reduction of endogenous PUM *in vivo*.

**Figure 6.**
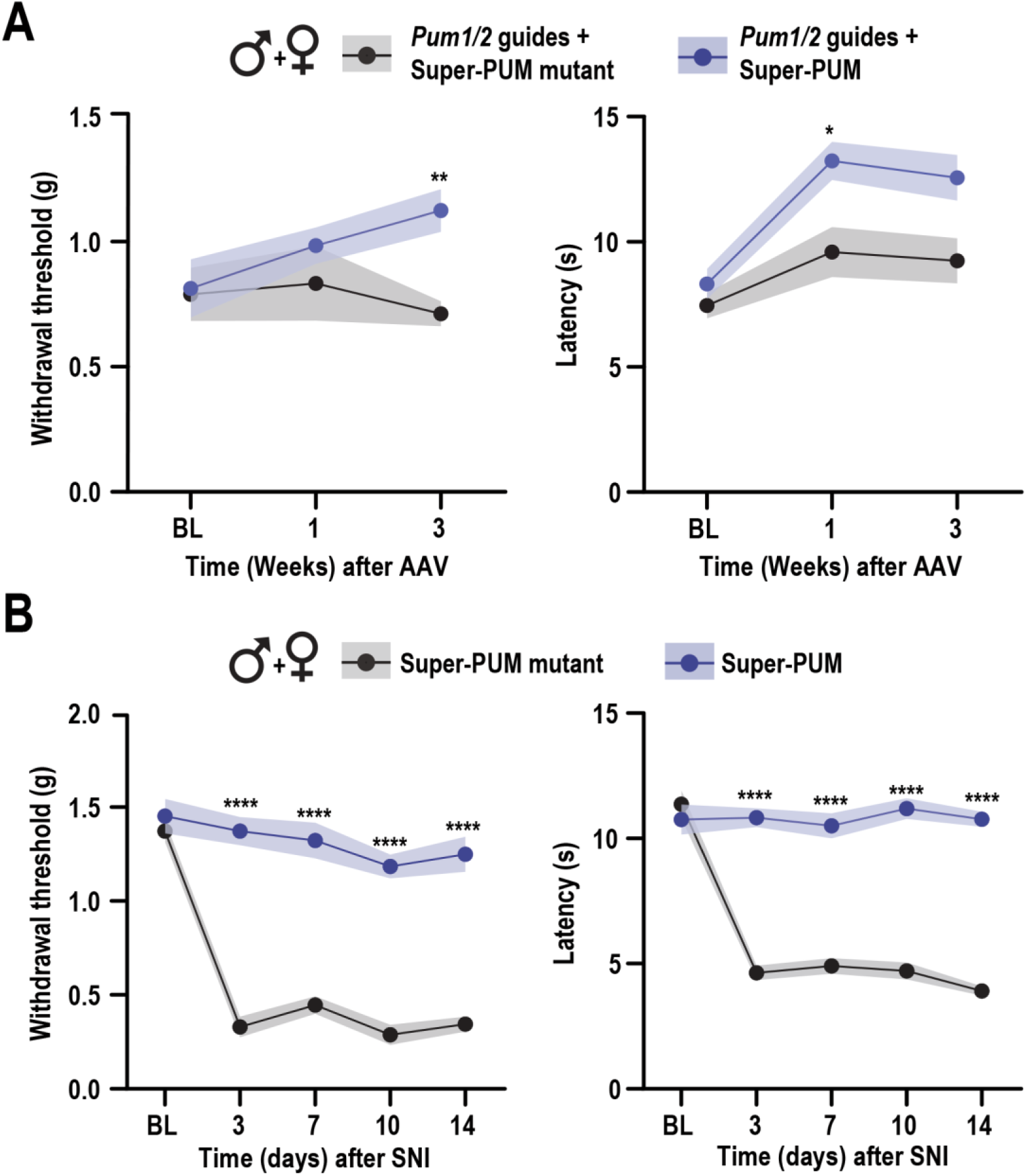
Super-PUM diminishes neuropathic pain following SNI. (**A**) Rescue of sensory abnormalities induced by PUM deletion through ectopic expression of super-PUM. Cas9 knock-in mice received *Pum1/2*-targeting guides together with AAV-super-PUM or AAV-super-PUM mutant. Mechanical withdrawal thresholds and thermal withdrawal latencies were assessed at baseline (BL), one week, and three weeks after viral delivery. Expression of super-PUM restored sensory thresholds in mice lacking PUM1/2, whereas the super PUM-mutant failed to rescue this phenotype. Data are presented as mean ± SEM (shaded area) of withdrawal threshold (g) or latency (s) from 6-7 mice per group. *P < 0.05 and **P < 0.01 were determined using two-way ANOVA followed by Bonferronís post hoc test. (**B**) Effects of AAVs on SNI-induced neuropathic pain. Super-PUM significantly attenuated the development of mechanical allodynia and thermal hypersensitivity following nerve injury compared to the super-PUM mutant. Data are presented as mean ± SEM (shaded area) of withdrawal threshold (g) or latency (s) from 10 mice per group. Measurements were taken at baseline (BL) and on days 3, 7, 10, and 14 after SNI in mice expressing either super-PUM or the super-PUM mutant. ****P < 0.0001 was determined by two-way repeated measures ANOVA followed by Bonferroni’s post hoc test.

To test if super-PUM can diminish injury-induced mechanical and thermal hypersensitivity, we leveraged the SNI model of neuropathic pain (*35*). SNI consists of ligation of the tibial and common peroneal branches of the sciatic nerve while sparing the sural branch, resulting in persistent mechanical and thermal hypersensitivity on the lateral surface of the paw. Mice were injected with AAV encoding either super-PUM or super-PUM mutant and subjected to SNI. In mice expressing the mutant construct, SNI produced robust mechanical allodynia and reduced thermal withdrawal latencies (**Fig. 6B**). In contrast, expression of super-PUM prevented the development of mechanical and thermal hypersensitivity. Together, these data demonstrate that enhancing PUM-dependent repression in sensory neurons is sufficient to suppress the development of characteristic neuropathic pain behaviors.

## DISCUSSION

Our data support four major conclusions. First, PUM1 and PUM2 are functionally redundant in maintaining normal mechanical sensitivity. Second, *Scn10a* is a functionally relevant target of PUM. Third, PUM regulates a network of VGSCs in the DRG. Fourth and finally, optimization of PUM repression with a synthetic chimera prevents the development of neuropathic pain. We discuss each finding below.

Genetic disruption of either PUM1 or PUM2 had minimal effects on baseline mechanical or thermal sensitivity. This contrasts with a strong phenotype observed following depletion of both PUM proteins. This observation is reminiscent of observations of redundancy among PUM homologues spread across the tree of life. For example, in *C. elegans*, *fem-3* binding factor-1 (FBF-1) and FBF-2 are required for progression through mitosis in the gonad (*36*). In mice, PUM1 and PUM2 are required for spermatogenesis (*37*). Additionally, PUM double mutants are not viable beyond embryonic day 8.5 (*38*). These findings raise an interesting question – why is redundancy so common in post-transcriptional gene control by PUM and its homologues? A possible explanation relates to robustness. One manner to preserve critical functions is through the maintenance of backup systems. We speculate that maintenance of two paralogous copies of PUM ensures that loss of either PUM1 or PUM2 due to mutational processes in DRG neurons does not compromise pain signaling, which is critically important for organismal survival (*39*).

Interestingly, the behavioral effect of PUM deletion differed between sensory modalities and inflammatory conditions. Combined loss of PUM increased baseline mechanical sensitivity without affecting thermal sensitivity, whereas PGE_2_ administration revealed enhanced heat hyperalgesia. These findings raise the possibility that increased expression of Na_v_1.7, Na_v_1.8 and Na_v_1.9 downstream of PUM deletion predisposes nociceptors to heightened inflammatory sensitization. Consistent with this hypothesis, PGE_2_ enhances Na_v_1.8 trafficking and sodium currents, while Na_v_1.9 has been shown to contribute substantially to PGE_2_-induced thermal hyperalgesia (*40, 41*). An unresolved but important question is if either inflammation or peripheral nerve injury alters PUM abundance or activity.

Na_v_1.8 plays a critical role in pain signaling and is a *bona fide* target of PUM. This finding contributes to a growing appreciation of its multimodal regulation. *Scn10a* is subject to meticulous regulation through transcriptional control, alternative splicing, and alternative polyadenylation (*42, 43*). Additional regulation is conferred through post-translational modifications on Na_v_1.8, protein partnerships that govern stability and localization (*41, 44, 45*). Our work reveals an additional layer of complexity in how this critical factor is regulated on a post-transcriptional basis. Broadly speaking, the identification of functionally important regulatory interactions between RNA-binding proteins and cis-acting elements in their targets remains technically challenging. In the context of pain, the sole genetically validated example is regulation of the non-selective sodium leak channel *Nalcn* by HnRNP L (*46*). This work made use of transgenic mice that lack PREs to exclude Scn9a/Na_v_1.7 as a relevant target. An important question moving forward is whether or not the PREs present in the *Scn10a* 3’UTR are necessary to confer regulation by PUM.

The approval of suzetrigine for acute post-surgical pain underscores the clinical importance of Na_v_1.8. However, it is unclear if blockade of Na_v_1.8 will be useful for the attenuation of neuropathic pain. Human sensory neurons exposed to suzetrigine remain able to generate action potentials, due to compensation by other VGSCs (*47*). This redundancy may underlie the incomplete analgesia observed clinically. A potential way to coordinately control multiple VGSCs is through synthetic post-transcriptional regulation. Here, we demonstrate that super-PUM reduces Na_v_1.8 abundance and function in sensory neurons and prevents neuropathic hypersensitivity following SNI. Given that PUM2 also associates with *Scn9a* and *Scn11a* transcripts, super-PUM may exert broader regulatory effects on sodium channel expression. Together, these findings support engineered PUM-mediated repression as a potential strategy for modulating nociceptor excitability and alleviating neuropathic pain.

We describe a novel protein containing the PUM RNA-binding domain fused to a highly efficient repressor. This creation of a compact fusion enabled heterologous repression using an AAV, as AAVs are size limited. Our strategy rests on the intrinsic specificity of PUM for RNA imparted through the modular repeats present in the C-terminal RNA-binding region. The general strategy of linking PUM RNA-binding domains to effectors has been leveraged in other contexts. For example, fusion of FBF-2 to a poly(A) polymerase promotes poly(A) addition of target substrates in *Xenopus* oocytes (*48*). An array of different effectors spanning splicing factors and translation activators diversify the use of this system for varied applications in synthetic biology (*49–51*). This work capitalizes on decades of fundamental research as a means to ameliorate neuropathic pain in a relevant preclinical model. Given that rare mutations in PUM are associated with developmental disability and ataxia, it is conceivable that this strategy will have salubrious applications that extend beyond pain (*52*).

To summarize, post-transcriptional regulation is an emerging mechanism in pain-associated plasticity. Many of these mechanisms hinge on the activity of a subset of RNA-binding proteins. Through the use of genome engineering, we discovered that PUM regulates *Scn10a*. Using synthetic biology, we showed that ectopic amplification of *Scn10a* repression renders mice resistant to neuropathic pain. The general strategy of exploiting information conferred by separable RNA-targeting and regulatory domains provides a range of possibilities for tailored post-transcriptional control.

## MATERIALS AND METHODS

### Mice

Genome editing was conducted using Rosa26-LSL-Cas9 knock-in (Cas9^lsl/+^) mice obtained from Jackson Laboratory (Stock No. 024857) (*53*). *Scn10a^Cre/+^* (Stock No. 036564) mice were used to generate the conditional Cas9 knock-in model (*54*). *Scn9a^PRE-Mut^* mice were genetically engineered by Cyagen Biosciences. In brief, Pumilio response elements were identified based on comparison to the defined PUM consensus motif (UGUANAUA) (*8, 11, 27*). The UGU elements were replaced with ACA to abrogate binding for each of the PREs (*9, 28*). Wild-type C57BL/6J mice were bred in our laboratory (Clinical Science Center, UW-Madison). Male and female mice aged between 8-12 weeks old were randomly assigned experimental groups. The individual groups were constrained to a maximum of three mice from the same litter of the same cage. All mice were housed under a 12-hour light/dark cycle with controlled ambient temperature (20– 26°C) and humidity (30–70%). Food and water were provided *ad libitum*. All animal procedures were conducted in accordance with approved protocols from the Institutional Animal Care and Use Committee (IACUC) at the University of Wisconsin-Madison (Protocol ID: M006582-A07).

### Behavioral assays

Mechanical hypersensitivity was assessed using calibrated von Frey filaments (BIOSEB Instruments BIO-VF-M)) to determine the paw withdrawal threshold in response to mechanical stimuli. Mice were placed in acrylic boxes with wire mesh floors and allowed to acclimate for 1 hour. Subsequently, von Frey filaments were applied to the plantar surface of the left hind paw for 2 seconds, and the up–down method was performed to calculate the mechanical withdrawal threshold in grams (*55*).

To assess heat sensitivity, the Hargreaves test was employed (*56*). Mice were habituated to the room 30 minutes before the test. Animals were placed on a hot plate maintained at 50°C and the time taken to observe nocifensive behavior was recorded. Nocifensive behaviors were considered when mice showed hind paw withdrawal or licking, stamping, leaning posture, or jumping. The Hargreaves test was performed to measure paw withdrawal latency in response to a focused radiant heat source. Mice were placed on a glass floor within an enclosed chamber, and a high-intensity infrared light beam was directed at the plantar surface of the left hind paw. The light intensity was set at 25-30% of maximum output, as per the manufacturer’s guidelines (IITC Model 390). Paw withdrawal latency was recorded in triplicate, with a minimum interval of 10 minutes between trials to minimize experimental variability. The cutoff time was 20 seconds for both thermal tests.

Prostaglandin E2 (PGE2) was used to induce inflammatory hypersensitivity. Mice received an intraplantar injection of 100 ng PGE_2_ (20 µL of 5 ng/µL in isotonic saline; Cayman Chemical, CAS 363-24-6) into the left hindpaw. Mechanical and thermal sensitivity were assessed at 3 hours and 1, 3, and 7 days following injection. For Na_v_ channel inhibitor studies, mice received intraplantar delivery of the Na_v_1.8 inhibitor suzetrigine (1 µg in 20 µL, MedChem Express, HY-148800) or the Na_v_1.7 inhibitor PF-05198007 (1 µg in 20 µL, Medchem Expess, HY-12883A) immediately after baseline behavioral testing.

### Spared nerve injury

Mice were placed on a heating pad and anesthetized with 3% isoflurane in oxygen. The lateral surface of the left thigh was shaved and disinfected with povidone-iodine (Med Vet International 50-252-2193) and 70% ethanol. A small incision was made through the biceps femoris muscle to expose the sciatic nerve and its three terminal branches: the sural, common peroneal, and tibial nerves. The common peroneal and tibial nerves were carefully ligated with 6-0 silk sutures (Med Vet International 50-252-2193) and transected distal to the ligation (approximately 1–2 mm), and a small segment of each nerve was removed to prevent reinnervation. Care was taken to avoid manipulation or damage to the sural nerve, which was left intact. Following surgery, mice were allowed to recover on a heated surface and were monitored until fully ambulatory. Mechanical allodynia and thermal hyperalgesia were performed at 3, 7, 10, and 14 days after surgery.

### Immunohistochemistry

Mice were briefly anesthetized with isoflurane and transcardially perfused with ice-cold phosphate buffered saline (PBS), pH 7.4 (Thermo Scientific J75889-K8) followed by 4% paraformaldehyde (PFA, Sigma-Aldrich 158127) in PBS. Lumbar DRG were collected and post-fixed overnight at 4°C with 4% PFA. Then, tissues were cryoprotected in 30% sucrose (Invitrogen 15503-022) for 3 days at 4°C. DRG samples were embedded in Tissue-Tek OCT compound (Sakura Finetek 4583) and 14 µm sections were collected. Tissues sections were washed twice with tris-buffered saline (TBS) (Thermo Scientific J60764.K7) and incubated with 10 mM sodium citrate (Fischer Scientific S279), 0.05% Tween 20 (Research Products International P20370), pH 6 for 1 hour. Sections were blocked with 10% goat serum (R&D Systems S13150), 0.3% Triton X-100 (VWR 0694), in TBS for 2 hours at room temperature using a humidified chamber. For staining involving mouse primary antibodies, goat anti-mouse Fab fragment (1:1,000, 115–001-003, Jackson ImmunoResearch) was included in blocking buffer. Tissues were incubated overnight at 4°C in primary antibodies diluted in incubation solution (1% goat serum, 0.3% Triton X-100, in TBS). Antoibody dilutions were as follows: rabbit anti-PUM1 (1:100, GeneTex GTX115280), rabbit anti-PUM2 (1:100, Bethyl A300-202A), guinea pig anti-TRPV1 (1:200, Neuromics GP14100), chicken anti-GFP (1:1000, Novus Biologicals NBP1-05423), mouse anti-Na_v_1.8 (1:1,000, NeuroMab), rabbit anti-Na_v_1.7 (1:200, Alomone Labs ASC-008) antibodies. Sections were washed three times with 0.1%Triton X-100 in TBS and incubated with the corresponding secondary antibody (1:1000, in incubation solution) or isolectin GS-IB4 (1:500, Vector Labs DL-1208) for 1 hour at room temperature. Sections were then washed three times with 0.1% Triton X-100 in TBS before cover slips were mounted using ProLong™ Glass Antifade (ThermoFisher Scientific P36984).

### Microscopy and image analysis

Images were captured on a Leica SP8 laser scanning confocal microscope with LAS X Life Science software. For experiments using the super-PUM mice and Na_v_1.8 staining, a Nikon TI2 Eclipse inverted light microscope with Nikon Elements software was used. Lumbar DRG sections were imaged using the respective 20x objective. Confocal imaging was performed with a pinhole of 1 Airy and scan speed of 400 Hz, using sequential acquisition to minimize spectral overlap. Data were collected from 4 or 5 mice per experimental condition. Images presented are representative projections of Z-stacks obtained from at least three DRG sections from the same sample. The number of PUM1 or PUM2 positive cells that colocalized with nociceptive neurons markers was divided by the total number of TRPV1 or IB4 positive cells and multiplied by 100 to calculate the percentage of positive cells. Fluorescence intensity was quantified using ImageJ by calculating corrected total cell fluorescence (CTCF) values. Regions of interest were drawn to measure the PUM1, PUM2, Na_V_1.7 or Na_v_1.8 fluorescence intensity specifically in GFP, TRPV1 or IB4 positive cells, according to the experiment. Results were calculated as either the mean of CTCF values or percent of at least 200 positive cells from a mouse sample and then, the mean of the 4-5 mice per experimental condition.

### RNA immunoprecipitation and real-time quantitative PCR (RIP-qPCR)

DRGs were collected in RNA immunoprecipitation lysis buffer - 50 mM Tris (Research Products International TG0040), pH = 8; 1 mM EDTA (Research Products International E14000); 2 mM MgCl_2_ (Research Products International M24000); 150 mM NaCl (Research Products International S23020); 0.5% NP-40 (Research Products International N59000) supplemented with 150 U/mL SUPERase In RNase inhibitor (Invitrogen AM2694) and 0.1 tablet/mL Pierce EDTA-free protease and phosphatase inhibitor (Thermo Fisher Scientific A32961). Each replicate consisted of all DRG tissue from 2 mice. Tissue samples were homogenized and centrifuged at 12,000 × *g* for 20 minutes at 4°C. A small amount of supernatant was reserved as input sample. The remaining supernatant was divided equally for immunoaffinity purification. 250 µL of supernatant was incubated with 10 µg of rabbit anti-PUM2 (Bethyl A300-202A) or anti-rabbit IgG (Proteintech 30000-0-AP) antibodies overnight at 4°C while rotating. The next day, 20 µl of protein A agarose beads (Cell Signaling Technology 9863) were added to the mixture and incubated for 3 hours at 4°C while rotating. The mixture was centrifugated at 2000 × *g* for 20 minutes at 4°C and washed 3 times with wash buffer (50 mM Tris, pH = 8; 1 mM EDTA; 2 mM MgCl_2_; 500 mM NaCl; 0.05% NP-40). For qPCR normalization, 10 ng of firefly luciferase (fLuc) mRNA (Promega L456A) was added to each sample. RNA was isolated by briefly vortexing the beads in 1 mL of Trizol (Invitrogen 15596018) and incubating at room-temperature for 5 minutes. 0.2 mL of chloroform (Sigma-Aldrich 34854) was added and vortexed for 10 seconds. Samples were centrifuged at 12,000 x g for 15 minutes at 4°C. mRNA was isolated from the aqueous phase by isopropanol precipitation.

cDNA was generated using 0.2 µg of total RNA by reverse transcription with the SuperScript VILO cDNA synthesis kit (Invitrogen 11754250), following the manufacturer’s guidelines. RT-qPCR was performed with iTaq Universal SYBR Green One-Step kit (Bio-Rad 1725121) on an Applied Biosystems Real Time PCR StepOne Plus machine in a total volume of 10 μL with 10 ng cDNA template. Cycling conditions were: initial denaturation at 95°C for 10 min, followed by 40 cycles of 95°C for 20 seconds, and 60°C for 1 minute, with fluorescence read after each cycle. Primer sequences are provided in Suppl. Table 1. All reactions were performed in duplicate. The mean Cт value for genes of interest were normalized to fLuc in the same sample. Then, values obtained from samples treated with rabbit anti-PUM2 or anti-rabbit IgG antibodies were used to calculate relative enrichment using the 2^-ΔΔCт^ method.

### Immunoblots

Lumbar DRGs were homogenized in ice-cold radioimmunoprecipitation buffer supplemented with 0.1 tablet/mL Pierce EDTA-free protease and phosphatase inhibitor Homogenates were centrifuged at 4°C for 20 minutes at 12,000 × g. Protein concentration in the supernatant was quantified by Pierce BCA protein assay (Thermo Fisher Scientific 23225). 40 μg of total protein was denatured at 95°C for 5 minutes in 1x Laemmli Sample Buffer (Bio-Rad 1610747) with β-mercaptoethanol (Fisher Scientific O3446I) before separation by 10% SDS-polyacrylamide gel electrophoresis and transferred to polyvinylidene difluoride membranes (Millipore IPVH00010). Then, membranes were blocked with 5% nonfat dry milk (Research Products International M17200) in TBS supplemented with 0.1% Tween-20 (TBST) for 1 hour. Membranes were incubated overnight at 4°C in TBST containing primary antibodies at the following concentrations: rabbit anti-PUM2 (1:1000, Bethyl A300-202A), rabbit anti-Na_v_1.7 (1:500, Alomone Labs ASC-008), mouse anti-Na_v_1.8 (1:500, NeuroMab 75-166), rabbit anti-Na_v_1.9 (1:500, Alomone Labs ASC-017), or rabbit anti-GAPDH (1:20,000, Proteintech 10494IAP). Membranes were washed three times with 1% milk in TBST prior to incubation with the corresponding anti-rabbit (1:10000, Jackson ImmunoResearch 111-036-144) or anti-mouse (1:5000, Jackson ImmunoResearch 115-036-062) horseradish peroxidase (HRP)-conjugated secondary antibodies for 1 hour at room temperature. HRP signal detection was achieved with the enhanced chemiluminescence system (Millipore WBKLS0500) according to the manufacturer’s instructions. GAPDH was used as an internal loading control to normalize protein expression levels. Full sized blots are provided in the supplement (**Suppl. Fig. 6)**.

### Guide RNA Synthesis and Delivery

Guide RNAs were generated as previously described (*26*). Briefly, *Pum1* and *Pum2* specific protospacer sequences were identified with CHOPCHOP using the GRCm38 genome assembly with the CRISPR/Cas9 and knockout settings (*57*). DNA oligos used for the generation of transcription templates are detailed in Suppl. Table 1. After *in vitro* transcription, 5’ triphosphate removal (Quick CIP, NEB M0525S), and RNA purification (RNA Clean & Concentrator-5, Zymo Research R1014), individual guide RNAs were pooled in equal amounts. Three individual guide RNA were delivered per target. Guide RNA pools were diluted 2:5 in *in vivo-*jetRNA (Polyplus 101000122) reagent to form RNA/polymer complexes (3 µg guideRNA/5 µL final concentration). 5 µL of RNA/polymer complexes were injected intrathecally into anesthetized animals at the L4/L5 vertebrae using a 31-gauge needle. Two injections were conducted with a one-week recovery period between injections.

### Cell Culture

Human embryonic kidney cells (HEK293T, ATCC) and mouse embryonic fibroblasts (NIH-3T3, ATCC) were maintained at 37°C in a humidified incubator with 5% CO_2_. Cells were cultured in Dulbecco’s Modified Eagle Medium (DMEM, Gibco 11995-065) supplemented with 10% fetal bovine serum (Corning 35-011-CVAll). Cell culture procedures were performed under sterile conditions in a biosafety cabinet.

### AAV Production and Delivery

AAV genome plasmids containing transgenes flanked by AAV inverted terminal repeats were generated using Gibson assembly. Plasmids encoding for wild-type, human PUM2 RNA binding domain (residues 705-1039) or an RNA-binding defective mutant (N923A, Y924A, Q927A, S959A, N960A, E963A) in the pcDNA3.1(+) backbone were generated by Gene Universal. All primer sequences are provided in Suppl. Table 1. Protein coding sequences were amplified with Q5 polymerase (NEB M0491S) with primers designed to introduce 30-40 bp overlapping, homologous regions between adjacent fragments. Murine CNOT7 coding sequence was amplified from cDNA from NIH-3T3 cells using Q5 polymerase. A P2A sequence was amplified from Addgene plasmid #187950. The pAAV-hSyn-DIO-EGFP backbone (Addgene plasmid #50457) was prepared via NcoI digestion (NEB R31933S). PCR products and vector backbone were combined with 2x HiFi Assembly mix (NEB M0541S) following manufacturer’s instructions.

To produce AAV particles, 15 cm plates of 70-80% confluent HEK293T cells were transfected with 5.7 µg AAV genome plasmid, 11.4 µg pHelper (Novapro V005569), and 22.9 µg pUCmini-MaCPNS1 (Addgene plasmid #185136) (40 µg total, 1:4:2 ratio). Transfections were performed with a 3.75:1 linear, MW 25,000 PEI:DNA ratio (Kyfora Bio 23966). Immediately before transfecting, media was changed to 5% FBS,1x Non-essential Amino Acids (Gibco 11140076) in DMEM. Media was exchanged at 6 and 64 hours after transfection. AAV particles were collected on the fifth day post-transfection. AAV particles were purified using the Takara AAVpro Purification Kit Maxi (All Serotypes) (Cat no. 6666), following kit instructions.

Real-time quantitative PCR was performed to quantify viral genome (vg) copy number. To destroy unencapsulated DNA, 2 µL purified AAV samples were brought to 100 µL in 0.05 U/mL DNase I (Thermo Scientific EN0521), 2 mM CaCl_2_ (Sigma-Aldrich C4901), 10 mM Tris-HCl (Research Products International T60050), 10 mM MgCl_2_. DNase I digestion was performed at 37°C for 1 hour. DNase I was inactivated by the addition of 25 mM EDTA and incubation at 70°C for 10 minutes. To liberate virally packaged DNA, 120 µL of 100 µg/mL Proteinase K (Roche 03115828001), 1 M NaCl, 1% w/v N-lauroylsarcosine sodium salt (Sigma-Aldrich L9150) solution was added to samples. Proteinase K digestion was carried out at 50°C for 2 hours followed by inactivation at 95°C for 10 minutes. Before qPCR, digested samples were diluted 1:300. Linearized AAV genome plasmid was used as standards for quantification from ∼1x10^2^-1x10^9^ ssDNA molecules/µL. vg per µL was calculated as the average concentration calculated from two qPCR reactions targeting two elements on the AAV genome plasmid (Table 1). iTaq Universal SYBR Green One-Step kit and an Applied Biosystems Real Time PCR StepOne Plus machine were used with the following cycling conditions: initial denaturation at 95°C for 10 min, followed by 40 cycles of 95°C for 20 seconds, and 60°C for 1 minute, with fluorescence read after each cycle. Melt curve analysis at the end of the run confirmed amplicon specificity.

For *in vivo* gene transduction, 1-3.5x10^11^ vg were injected intrathecally at the L4/L5 vertebrae and allowed to induce for two weeks.

### Super-PUM luciferase reporter assays

HEK293T cells were seeded at 5-10,000 cells per well in 100 μl of growth medium in a 96-well white-walled plate. To deplete endogenously expressed PUM, the next day, cells were transfected with 200 nM pool of two antisense oligonucleotides targeting *Pum1* and *Pum2* or a scrambled, non-targeting control (Suppl. Table 1) using Lipofectamine 3000 transfection reagent (Thermo Scientific L30000001). Plasmids encoding Nanoluciferase with a 3’UTR containing 3xPRE or 3x mutated PRE sequences were used to monitor PUM activity (*13*). The firefly luciferase plasmid pGL4.13 was used as the transfection efficiency control. Full-length PUM2 was expressed with pCF3F-Pum2, super-PUM was expressed with super-PUM or super-PUM mutant cloned into the MCS of pcDNA3.1(+) (*58*). 24 hours after ASO transfection, cells were transfected with a total of 100 ng DNA per well, consisting of 5 ng fLuc, 10 ng nLuc, and 85 ng of PUM plasmid, with Lipofectamine 3000. The transfected cells were incubated another 24 hours. Luciferase activity was measured using the Nano-Glo Dual-Luciferase Reporter Assay system (Promega N1610) and a BioTek Synergy H1 plate reader. Experiments were performed with eight total replicates. Statistical significance was assessed using an unpaired t test.

### Purification of PUM2 RBD

Expression and purification of the PUM2 RBD were adapted from previous protocols (*5*). His-tagged PUM2 RBD was expressed in Rosetta2 (DE3) *E. coli* using a pDEST527 vector. Cells were grown at 37°C to an OD of ∼0.6 before overnight induction with 0.2 mM IPTG at 16°C. Frozen cell pellets were resuspended in 50 mL lysis buffer (20 mM Tris-HCl pH 8.0, 500 mM NaCl, 20 mM imidazole, 5% glycerol, 0.5 mM TCEP) spiked with cOmplete EDTA-free protease inhibitor (Roche 11873580001), sonicated, and centrifuged. The supernatant was applied to 5 mL Ni-NTA resin (Qiagen 30230) pre-equilibrated with lysis buffer and incubated for approximately 1 hour. The resin was then washed with lysis buffer before equilibration with 2 CV of Low-Salt Buffer (20 mM Tris-HCl, pH 8.0, 50 mM NaCl, 20 mM imidazole, 0.5 mM TCEP) and subsequent elution with Elution Buffer (20 mM Tris-HCl, pH 8.0, 50 mM NaCl, 300 mM imidazole, 0.5 mM TCEP). The eluate was treated overnight at 4°C with in-house purified TEV protease and thereafter purified over a 5 mL Heparin HiTrap column (Cytiva 17040703) using linear gradient beginning with 95% Buffer A (20 mM Tris-HCl, pH 8.0, 0.5 mM TCEP) to increasing Buffer B (20 mM Tris-HCl, pH 8.0, 2M NaCl, 0.5 mM TCEP). Heparin fractions were pooled, concentrated, and additionally purified using a HiLoad 16/600 Superdex 75 pg column (Cytiva 28989333) in Filtration Buffer (20 mM Na-HEPES pH 7.4, 150 mM NaCl, 1 mM DTT, 10% glycerol). Peak gel filtration fractions were pooled, concentrated, and flash frozen in liquid nitrogen before storage at -80°C.

### Electrophoretic Mobility Shift Assay

Oligonucleotides containing putative PREs in SCN9A or SCN10A were ordered from IDT. These constructs contained 5 flanking nucleotides adjacent to the PRE alongside a 5′ FAM label. FAM-labelled RNAs were annealed in 5 nM reactions in RNA folding buffer (20 mM Na-HEPES pH 7.4, 150 mM NaCl, 1 mM DTT, 0.1% Tween-20) by incubating at 95°C for 5 minutes followed by incubation on ice for at least 15 minutes. After annealing, the samples were spiked with yeast tRNA (Invitrogen AM7119) to a final concentration of 200 µg/mL. Purified PUM2 RBD was serial diluted 2-fold beginning at 4 µM in RNA Folding Buffer lacking Tween-20 to yield 14 concentrations along with a no protein (buffer only) condition. 10 µL annealed RNA was mixed with 10 µL PUM2 RBD dilutions, yielding final binding reaction conditions of 20 mM Na-HEPES pH 7.4, 150 mM NaCl, 1 mM DTT, 0.05% tween-20, 100 ug/mL tRNA, 2.5 nM RNA, and protein concentrations beginning at 2 µM. The reactions were incubated at 25°C in a thermocycler for 2-4 hours before addition of 5 µL loading buffer (15% Ficoll 400 with 0.005% bromophenol blue). 10% TBE gels (Invitrogen EC62755BOX) were pre-run for at least 30 minutes in 1X TBE before loading 10 µL of the binding reactions to each well. Gels were run at 100V for 30-40 minutes and imaged on an Amersham Typhoon Imager (GE Healthcare). Band intensities were quantified in FIJI/ImageJ using the plot lanes tool, which facilitated calculation of fraction bound for each lane in the gels. K_d_’s were calculated in Graphpad Prism using a specific binding equation. Each EMSA was performed in three independent replicates, and *K*_d_’s are reported as average ± standard error of the mean (SEM).

### RNA structure prediction

RNA structure prediction was performed by using the Fold module in the RNAstructure webserver (*59*). All default parameters were used except that the temperature was set to 25°C.

### Electrophysiology

DRG neurons were cultured as previously described (*60*). The following day, whole-cell voltage-clamp recordings were obtained at room temperature (23°C) using a MultiClamp 700B amplifier and Clampex 11 software (Molecular Devices). Signals were digitized using a Digidata 1440A analog-to-digital converter, sampled at 20 kHz, and filtered at 0.3-3 kHz. Patch pipettes (internal resistance: 2-4 MΩ) were filled with (in mM): 140 CsF, 1 EGTA, 2.5 NaCl, 2.5 ATP magnesium salt, 20 HEPES. Cells were bathed in a solution containing (in mM) 130 NaCl, 20 TEA-Cl, 3 KCl, 1 MgCl_2_, 1 CaCl_2_, 10 HEPES, 10 D-glucose, 0.1 CdCl_2_. Series resistance was compensated by 60-80%. Cells were held at - 70 mV, and currents were evoked by 300 ms depolarizing steps from -70 to +25 mV in 5 mV increments. Residual capacitive transients were corrected offline by subtracting a linearly scaled subthreshold trace.

### Statistical analysis of results

All data were analyzed using GraphPad Prism software. For comparisons between two groups, unpaired t-test was applied. For comparisons involving two variables, two-way ANOVA analysis followed by Dunnett or Bonferroni’s post hoc tests were used. Results were presented as mean ± SEM. Statistical significance was set at P < 0.05. Sample sizes were determined based on previous publications in the field, which have consistently reported biologically and statistically meaningful results. During data acquisition and quantification analysis of all tissue samples and behavior data, the experimenter was blinded to the condition or treatment that samples or mice received.

## ACKNOWLEDGEMENT

This work was supported by NIH grants R01NS114018, R01NS147491, (Z.T.C.), 653 T32HL007936 (A.R.M.), and R01GM150468 (A.C.G.). This work was supported in part by the NIH, National Institute of Environmental Health Sciences (ZIA-ES050165, T.M.T.H.). The contributions of the NIH authors are considered Works of the United States Government. The findings and conclusions presented in this paper are those of the authors and do not necessarily reflect the views of the NIH or the U.S. Department of Health and Human Services

## AUTHOR CONTRIBUTIONS

Conceptualization: G.G., A.C.G., and Z.T.C.

Methodology: G.G., J.B.S., A.R.M., J.P.F., K.A.B., K.M., S.R., A.C.G., R.A.P., T.M.T.H, Z.T.C.

Investigation: G.G., J.B.S., A.R.M., J.P.F., K.A.B., K.M., S.R., A.C.G., R.A.P., T.M.T.H, Z.T.C.

Supervision: A.C.G., R.A.P., T.M.T.H, Z.T.C.

Writing, original draft: G.G. and Z.T.C.

Writing, review & editing: G.G., J.B.S., A.R.M., J.P.F., K.A.B., K.M., S.R., A.C.G., R.A.P., T.M.T.H, Z.T.C.

## AI DECLARATION

AI was not used in the preparation or editing of this manuscript.

