## Supplement for "Engineered post-transcriptional regulation of *Scn10a* by Pumilio ameliorates neuropathic pain in mice"

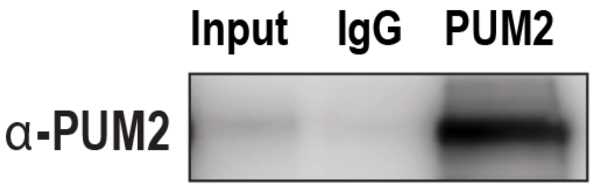


**Supplementary Figure 1.** Representative immunoblot showing efficient capture of PUM2 from input and immunoprecipitated fractions with IgG or PUM2 antibodies.

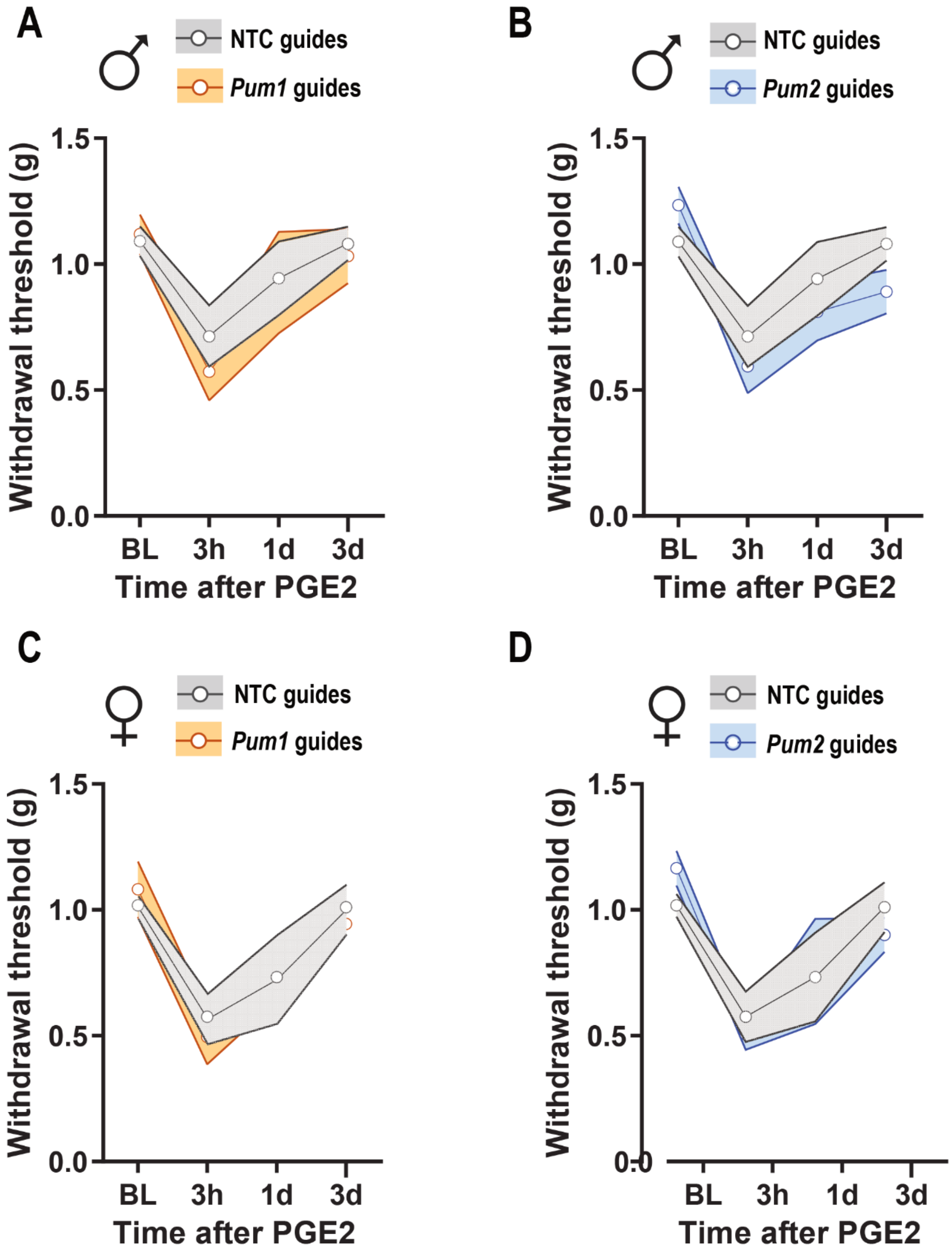


**Supplementary Figure 2.** Mechanical sensitivity in male (**A-B**) and female (**C-D**) Cas9 knock-in mice treated with RNA guides targeting *Pum1, Pum2*, or a nontargeting control (NTC) following intraplantar delivery of PGE2 (100 ng). Lines represent the mean ± SEM (shaded area) from 5-6 mice per group. Measurements were performed before PGE2 injection (baseline, BL) and at 3 hours (3h), 1 day (1d), and 3 days (3d) post-injection. Not significant differences were detected by two-way ANOVA.

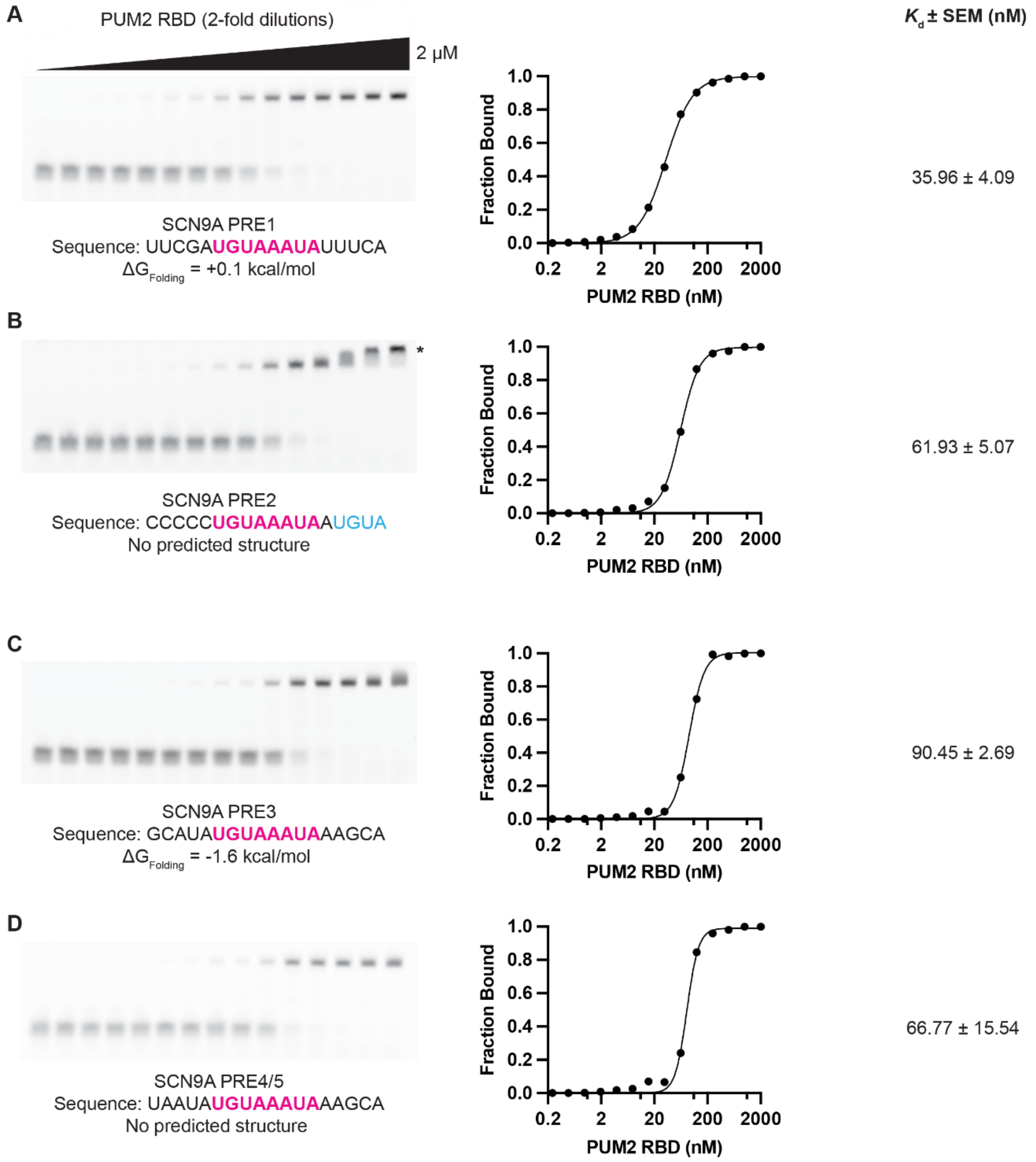


**Supplementary Figure 3.** PUM2 binds with Pumilio response elements (PRE) situated in the *Scn9a* 3′UTR. (**A-D**) Electrophoretic mobility shift assays showing binding of increasing concentrations of recombinant PUM2 RNA-binding domain to the five predicted PRE (PRE1–PRE5) identified within the canonical *Scn9a* 3′UTR. A secondary shift is indicated with an asterisk in panel B. Quantification of EMSA binding curves are provided on the right. Folding energies for RNA oligos were calculated in the RNAstructure Webserver [1].

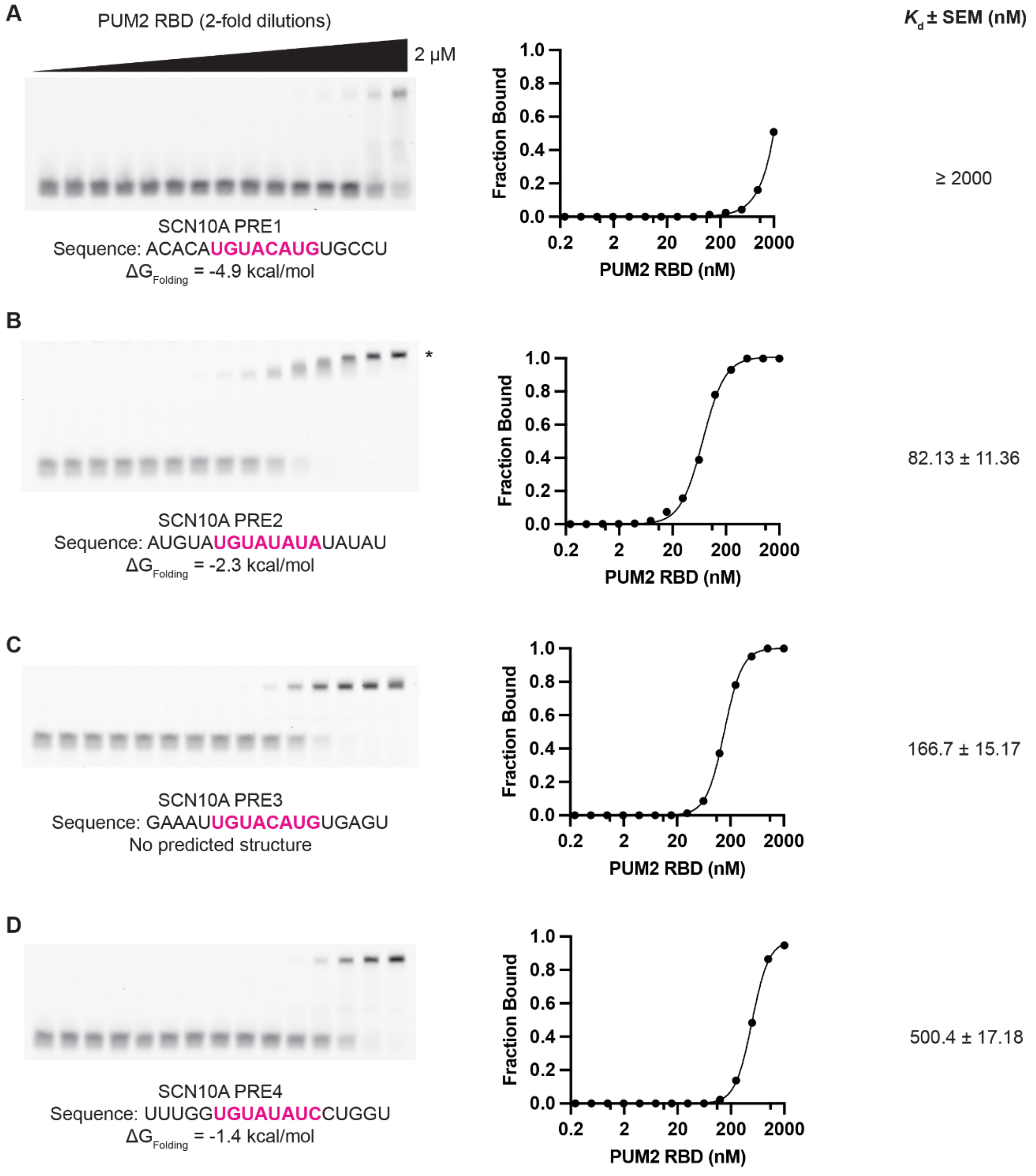


**Supplementary Figure 4.** PUM2 binds with Pumilio response elements (PRE) situated in the *Scn10a* 3′UTR. (**A-D**) EMSAs on four predicted PRE (PRE1–PRE4) in the *Scn10a* 3′UTR. A secondary shift is indicated with an asterisk in panel B. Quantification of EMSA binding curves are provided on the right. Folding energies for RNA oligos were calculated in the RNAstructure Webserver [1].

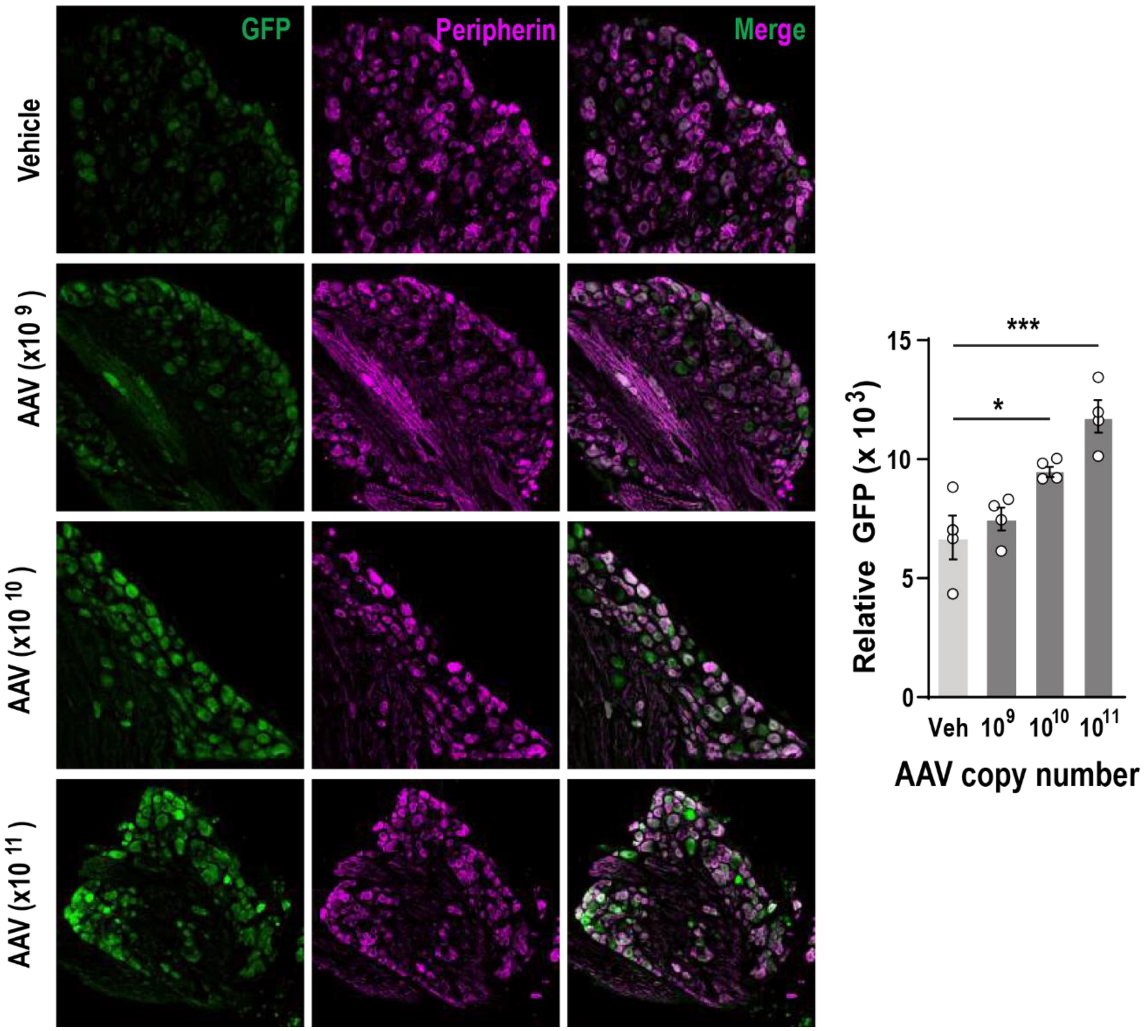


**Supplementary Figure 5.** Expression of AAV encoded GFP. AAVs encoding eGFP were injected intrathecally at the indicated viral genome copy numbers. Lumbar DRGs were harvested and stained for GFP and peripherin. Relative florescence quantification for GFP is plotted within the peripherin population. Bars represent the mean ± SEM of relative intensity values from n = 4 mice per group. *P < 0.05 and **P < 0.01 were determined by one-way ANOVA followed by Dunnett´s multiple comparisons test.

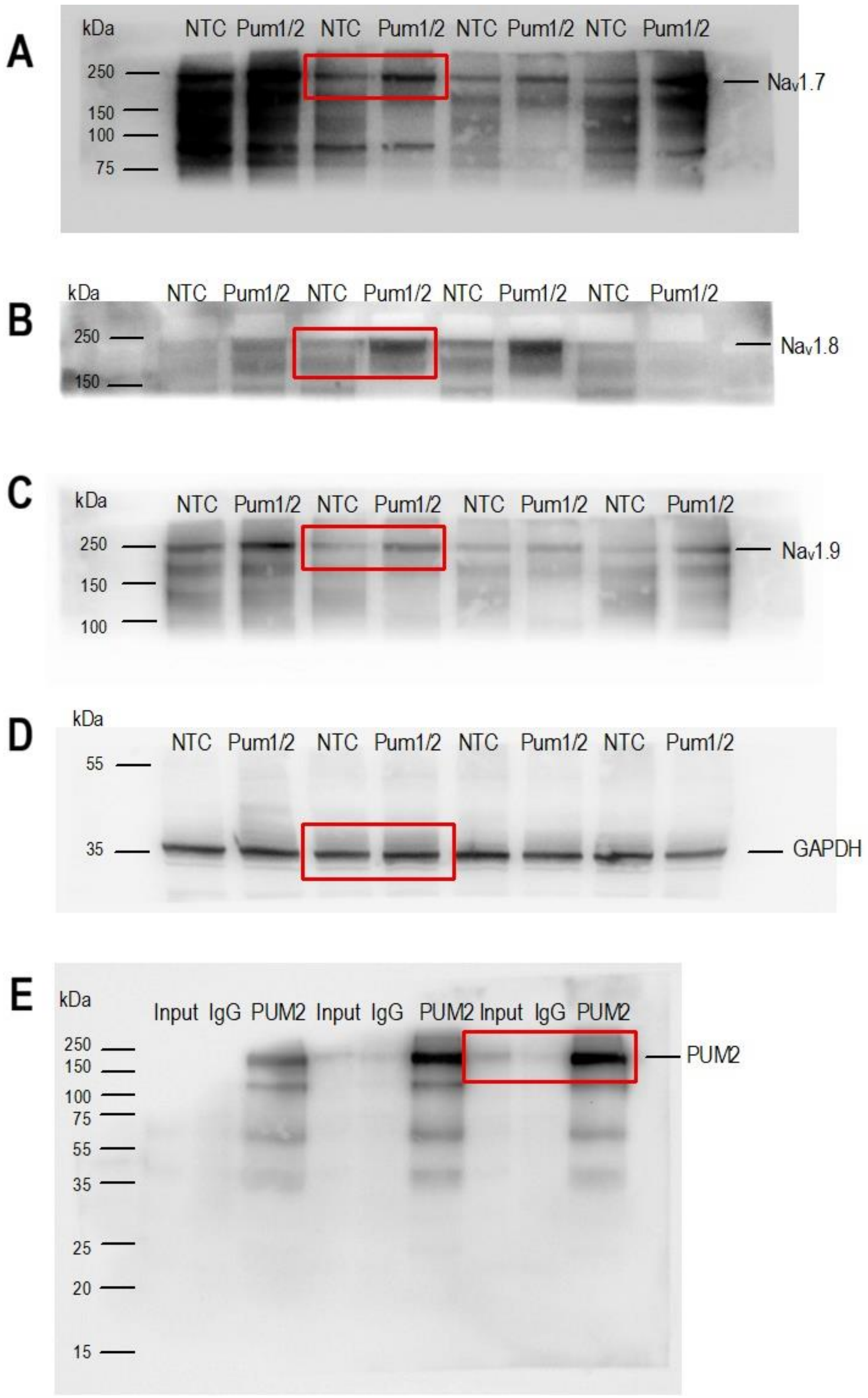


**Supplementary Figure 6.** Full-length Western blot membranes. Membranes corresponding to the immunoblots presented in Figure 2D (**A-D**) and Supplementary Figure 1 (**E**). Red boxes indicate the representative blots selected for presentation in the corresponding figures.
